# Extracellular glutamine is essential in cell division and flagellar function in bloodstream forms of *Trypanosoma brucei*

**DOI:** 10.1101/2025.11.17.688794

**Authors:** Flávia Silva Damasceno, Sabrina Marsiccobetre, Rodolpho Ornitz Oliveira Souza, Matheus Issa, Otavio Henrique Thiemann, Andrea Sorrentino, Katherine Figarella, Michael Duszenko, Ariel Mariano Silber

## Abstract

*Trypanosoma brucei* is the etiologic agent of African trypanosomiasis, also known as sleeping sickness. In the mammalian host, *T. brucei* is present as replicative and non-replicative bloodstream forms (BSF) referred to as long slender and stumpy, respectively. In the insect vector, the developmental forms are procyclic (PCF), epimastigote, and metacyclic trypomastigote. It is well established that BSF use glucose as the main energy source, and PCF can use glucose but, in its absence, uses amino acids, mainly proline. Due to these facts, the metabolism of amino acids has been well studied in PCF, but less attention has been given to it in BSF. In the present work, we unveil the participation of glutamine (Gln), the most abundant amino acid in the human blood, in BSF proliferation. According to the data presented herein, in the absence of extracellular Gln, cell cycle progress, as well as kinetoplast and flagella distribution and function among daughter cells, are widely compromised, yielding polyploid multiflagellated cells with reduced motility. In addition, the Gln-poor medium resulted in cells with a massive alteration in the proteome glutamylation. These alterations were reverted when Gln was added to the culture media. An analysis of the glutamylproteome of cells incubated in Gln-poor medium, followed by a Gene Ontology (GO) analysis, suggested a relationship between glutamylation and biological processes such as gene expression and regulation, cell cycle, and bioenergetics metabolism, in addition to cytoskeleton dynamics in BSF of *T. brucei*.

## Introduction

The protozoan parasite *T. brucei* is the causative agent of African Trypanosomiasis, commonly known as sleeping sickness or Human African Trypanosomiasis (HAT) in humans and Nagana in animals. This pathogen exists in various forms throughout its complex lifecycle, which involves an insect vector (the diptera Tsetse) and a mammalian host. Within the latter, *T. brucei* bloodstream form (BSF) circulates in blood, evading the immune system through sophisticated mechanisms known as antigenic variation [reviewed by (1)]. In the insect vector, the predominant form is the PCF, which invades the salivary glands to differentiate into the replicative epimastigote and finally the metacyclic trypomastigote. The metacyclic trypomastigote is the infective form inoculated into a mammalian host during the tsetse’s bloodmeal (1,2).

It is well established that in the absence of glucose, PCF have an elaborate energy metabolism based on amino acid consumption, mainly proline (2–4). In contrast, BSF utilizes glucose as the primary energy source, with a small amount of ATP generated in the mitochondria, using acetyl-CoA and/or threonine as substrates, depending on the cellular conditions (5,6). Consequently, limited attention has been given to the biological role of amino acids in this form. Among the relevant information available, aspartate is used as a precursor of pyrimidine synthesis (7), methionine is metabolized as an intermediate of the polyamine pathway by methyltransferases (8), while cysteine is an essential factor for BSF proliferation (9). In the specific case of Gln, it was described as indispensable for BSF proliferation *in vitro* (10). Additionally, a metabolomic analysis showed that Gln can feed mitochondrial pathways and act as the main donor of amino groups to the nitrogen pool of the parasite, participating in the synthesis of nucleotides, other amino acids, and other nitrogenous compounds (11).

The participation of amino acids in the post-translational modifications (PTMs) has been described in the PCF of *T. brucei*. In trypanosomatids, the cytoskeleton is composed of microtubules, which are responsible for the cell shape and constitute the mitotic spindle, the flagellar axoneme, the basal body of the flagellum, and the subpellicular corset. In general, the functional diversity of microtubules, central to a wide range of cellular processes, is partly driven by post-translational modifications. Among the best-characterized are acetylation, phosphorylation, polyglutamylation, polyglycylation, palmitoylation, polyamination, detyrosination, and glutamylation. It was demonstrated that in PCF the microtubules are extensively glutamylated (12–17) and that this is related to their participation in events such as positioning of organelles, intracellular transport, mitosis, cytokinesis, and cell motility (14,18,19).

The present work shows that the presence of Gln, the most abundant amino acid in human blood (i.e., the environment where the BSF develops), is critical for several seminal biological processes such as cytokinesis and motility. We demonstrate here that Gln induces protein glutamylation and polyglutamylation in *T. brucei* BSF. The analysis of the glutamyl-proteome and its alteration in parasites proliferating in the absence of Gln led us to conclude that glutamylation and polyglutamylation in BSF of *T. brucei* regulate several biological functions, such as gene expression and regulation, cell cycle progression, and bioenergetics metabolism, in addition to cytoskeleton dynamics.

## Results

### Extracellular glutamine, but not its internal biosynthesis, is critical for cell growth and cell cycle progression

Given the importance of Gln in BSF, the relevance of its uptake from the external medium versus its biosynthesis was investigated. As shown in **Fig. 1 A**, uptake of both Gln and Glu occurs in BSF in a saturable way and is about three times higher for Glu than for Gln, even though the Gln concentration in HMI-9 is much higher (584 mg L^-1^ versus 75 mg L^-1^). This led us to consider that Gln is produced intracellularly by the action of a Glutamine Synthetase (GS), an enzyme that forms Gln from Glu, ATP, and NADH. Since a putative GS (*Tb*927.7.4970), ortholog to the previously characterized GS of *T. cruzi* (20), is annotated in the *T. brucei* genome database, we used BSF cell extracts to measure the respective enzyme activity. In fact, GS activity was robustly detected even with as low as 10 µg of parasite lysate (**Fig. 1 B**).

**Figure 1.**
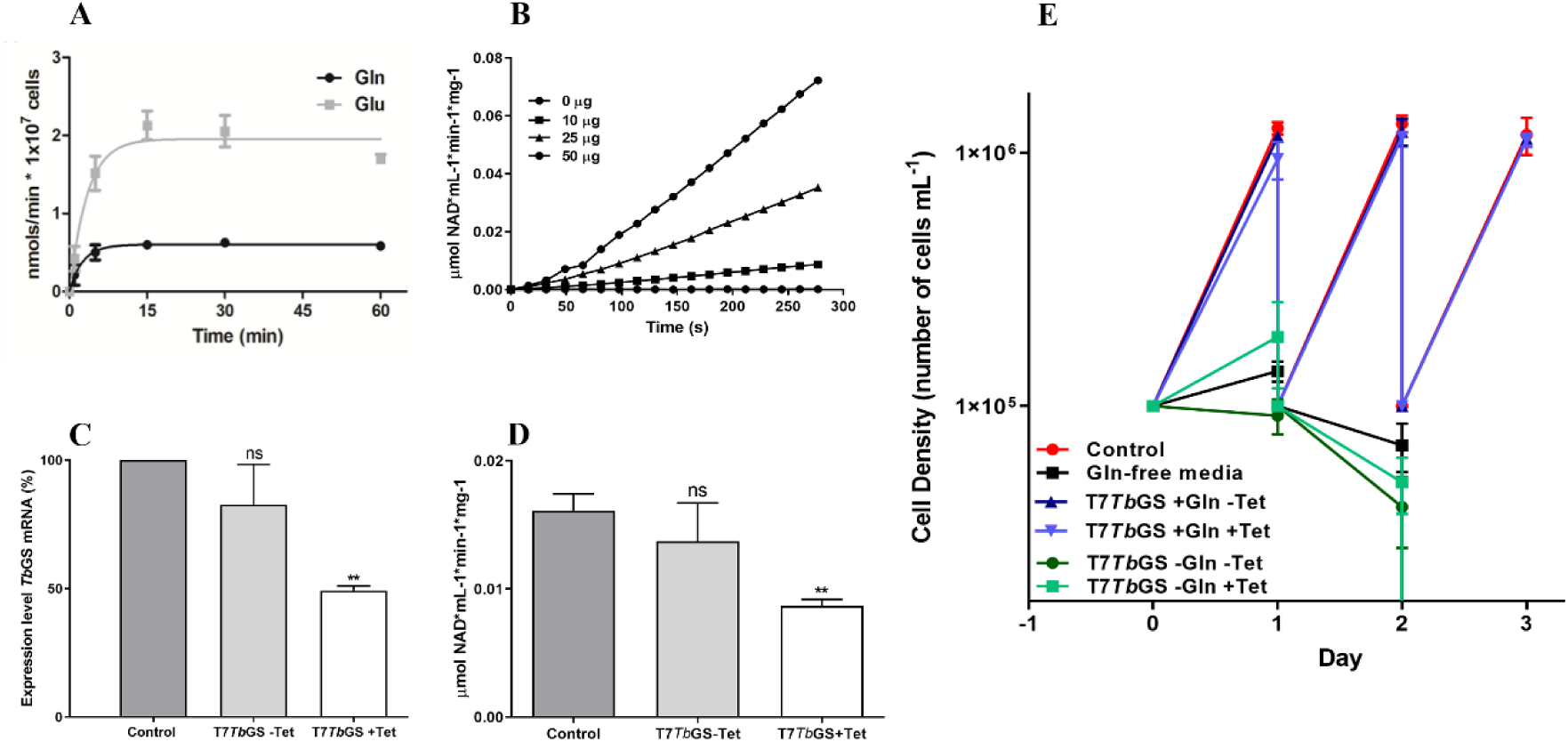
Glutamine and glutamate uptake, glutamine synthetase activity, and the effect of glutamine synthetase knockdown on BSF proliferation. (A) Time course of Gln and Glu uptake from the external medium. (B) Time course GS enzymatic activity using different amounts of BSF total protein extracts. (C) and (D) BSF of *T. brucei* were transfected with construct p2T7*Tb*GS, maintained in the HMI-9 medium and induced (+Tet) or not induced (-Tet) by the addition of tetracycline. After 48 h of cultivation, were evaluated (C) Transcripts for the TbGS mRNA were evaluated by qRT-PCR, and (D) TbGS activity in the parasite crude extracts. Statistical analysis: One-way ANOVA followed by Tukey’s test. P < 0.05. (E) Proliferation of *T. brucei* BSF: Not transfected (control) and transfected cells with the construction p2T7*Tb*GS were induced or not (+tet; -tet) by tetracycline addition in the presence or not of Gln (+Gln; -Gln). The culture was followed up by counting the cells in the Neubauer chamber over 3 days.

To study GS function in more detail, we transfected exponentially growing BSF parasites with the p2T7*Tb*GS construct and selected positive cells by the addition of phleomycin. The *Tb*GS knockdown was induced thereafter by adding tetracycline (2 µg mL^-1^) to the culture medium. In this way, expression of the *Tb*GS mRNA was reduced by 50.7% leading to a 54% decrease in GS activity in induced parasites compared to not transfected or uninduced parasites (**Fig. 1 C** and **Fig. 1 D**). However, cell proliferation of parasites was not altered although their GS activity was diminished by half (**Fig. 1 E**), when compared with non-induced parasites in HMI-9. This indicates that the supply of external Gln may compensate for the partial deficiency of Gln biosynthesis. Therefore, RNA_i_-induced (+tet) or non-induced (-tet) parasites were incubated in Gln-depleted HMI-9 to assess their ability to proliferate under this condition. Remarkably, both (Gln knockdown and control) parasites failed to proliferate (**Fig. 1 E**), demonstrating that extracellular Gln is critical to BSF cell proliferation, and that the intracellular production of Gln by the GS is unable to compensate for it.

As the cells were not affected by the GS knockdown but were affected by the depletion of extracellular Gln, we decided to investigate in more detail the participation of the extracellular Gln in cell proliferation and the cell cycle. To determine the critical concentration of external Gln for proliferation, we initially incubated the parasites in HMI-9 (control) or in a Gln-depleted HMI-9 in which we supplemented gradually decreasing concentrations of Gln (tenfold reduction of Gln every 24 h). The cells cultured in decreasing Gln concentration showed a normal proliferative profile, when compared to the control, until a concentration of 5 µg mL^-1^ of Gln. At a Gln concentration of 0.5 µg mL^-1^, the parasite’s proliferation was reversibly arrested, since supplementation of Gln to reestablish its original concentration led to the normal proliferation rate (**Fig. 2 A**).

**Figure 2.**
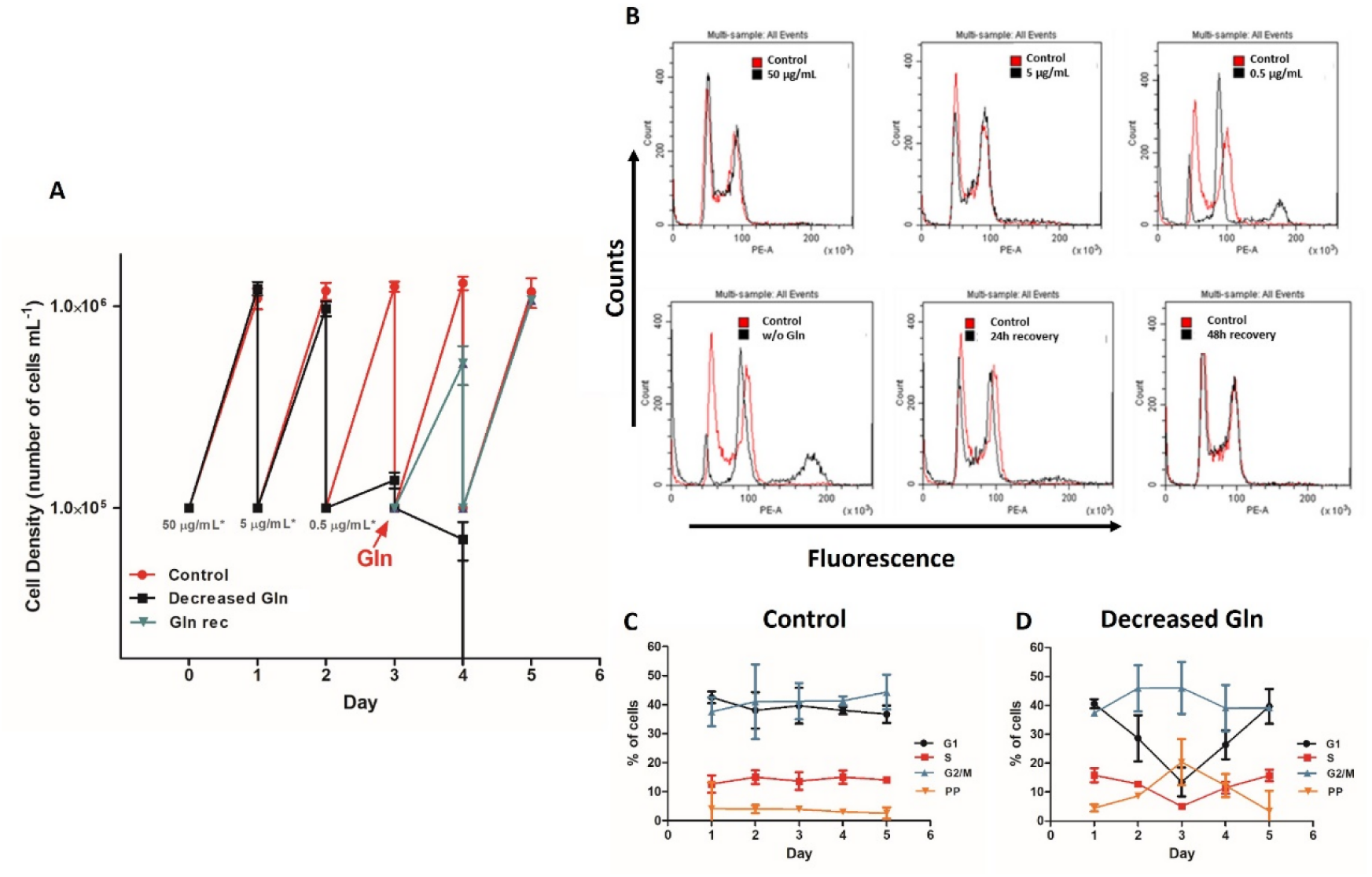
Effect of a gradual decrease in glutamine concentration on BSF proliferation and cell cycle. **(A)** Proliferation curves of *T. brucei* BSF. The parasites were maintained in a medium with a decreasing concentration of Gln each day (50, 5, 0.5 µg mL^-1^). After cultivation in the Gln 0.5 µg mL^-1^ media (third day), the parasites were recovered in complete Gln medium. The culture was followed up by 5 days, after daily counts, the cells were diluted back to their initial density (10^5^ mL^-1^), and the cell cycle was analyzed by flow cytometry: **(B)** Histogram overlay of daily analysis of cell cycle following the proliferation curve. **(C** and **D)** Representative graphs of three independent cell cycle cytometry analyses follow-up of the growth curve. **Control**: cells maintained in the complete HMI-9; **Decreased Gln**: cells kept in the gradually decreasing Gln medium.

Considering the effect of the extracellular Gln on proliferation, we were interested in analyzing the cell cycle of the parasites proliferating at different Gln concentrations. We observed that parasites kept in HMI-9 with decreasing Gln concentrations showed an altered distribution of the cell cycle phases when compared to those kept in HMI-9 (control). The cytometry profile observed in cells maintained in low Gln concentration showed an increased quantity of DNA, suggesting polyploidy (**Fig. 2 B, C,** and **D**). Interestingly, when the standard concentration of Gln was re-established in the medium, the cells recovered their proliferation rates and showed the same distribution of cell cycle phases, compared to the control (**Fig. 2B, C,** and **D**). These results demonstrate that although the parasites have an active GS, they are dependent on extracellular Gln to proliferate, and the low levels of this amino acid produce a failure in the BSF cell cycle. This indicates that Gln biosynthesis does not meet the cell’s requirement for this amino acid to maintain normal cell proliferation.

Replacement of Gln in the cell culture, after cultivation in Gln-free media, results in the recovery of proper proliferation and a normal cell cycle profile. To detect the minimal concentration of Gln that is necessary for maintaining the cells’ proliferation, the parasites were maintained for 24 h in Gln-free HMI-9 medium and then re-exposed to different concentrations of Gln. The recovery of the cells was Gln-dependent during the first 24 h (**Fig. 3 A**). After 48 h, the cells fully recovered their proliferation rates at the levels of the control when exposed to 584, 50, or 5 µg mL^-1^ but not 0.5 µg mL^-1^ Gln (**Fig. 3 A**).

**Figure 3.**
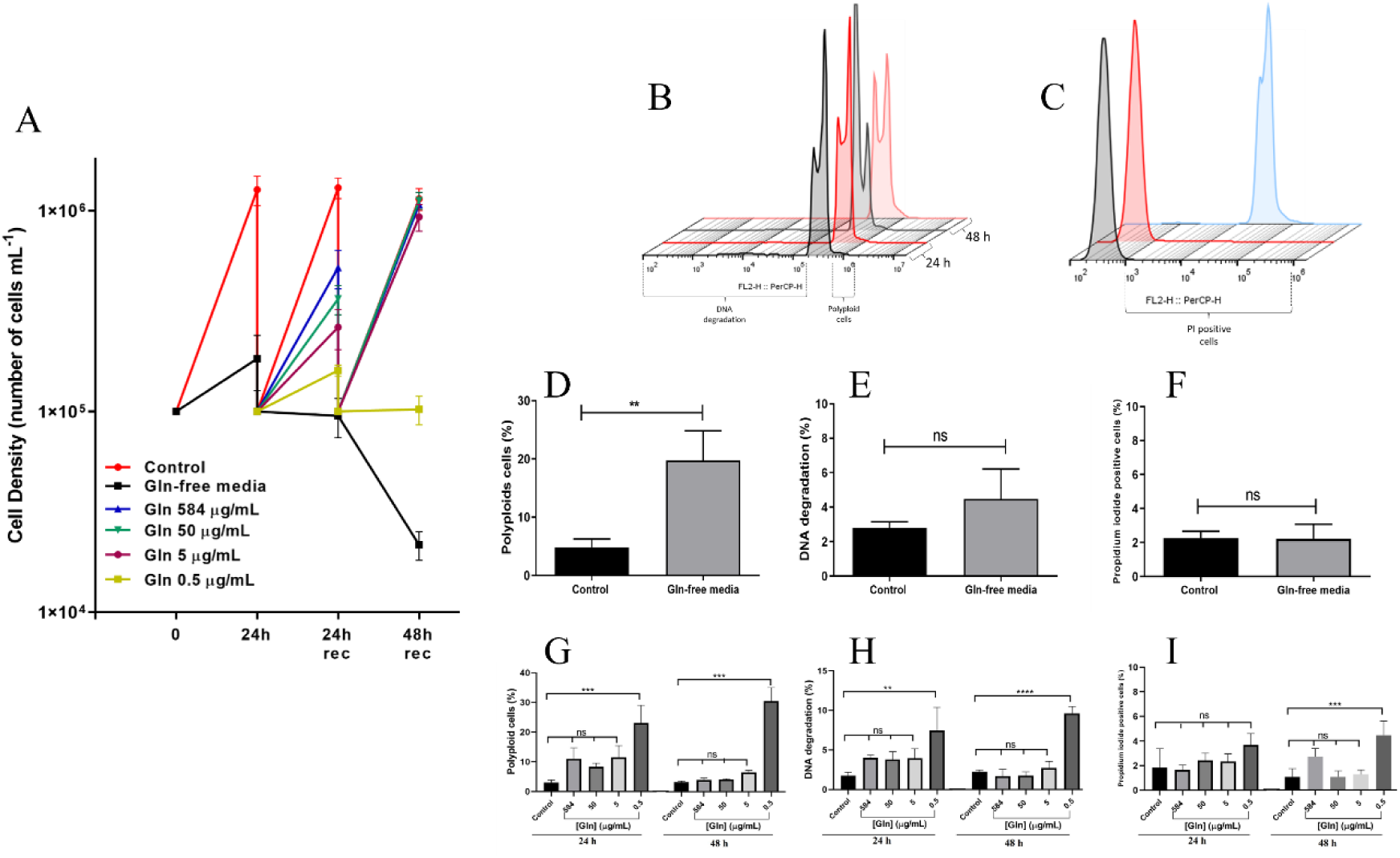
Bloodstream forms of *T. brucei* recovered in different concentrations of Gln. (**A**) Parasites were maintained in complete HMI-9 medium (control) or Gln-free HMI-9 medium for 24 h. The Gln-free culture was then split into five parts and recovered with different concentrations of Gln (584, 50, 5, and 0.5 µg mL^-1^). The cultures were monitored for three days; parasites were counted daily and diluted back to the initial concentration (10⁵ mL^-1^). Representative histograms (prepared with FlowJo version 10.10.0) show (**B**) cell cycle and (**C**) membrane integrity analyses by flow cytometry. Parasites maintained in complete HMI-9 medium are shown in black/gray, those in Gln-free medium in red, and permeabilized cells stained with propidium iodide (positive control) in blue. (**D–F**) To assess polyploidy, DNA degradation, and membrane integrity in BSF of *T. brucei*, cells were maintained in complete HMI-9 medium (control) or Gln-free medium for 24 h and analyzed for changes in cell cycle progression, DNA integrity, and membrane integrity. (**D**) Percentage of cells with DNA content suggestive of polyploidy; (**E**) percentage of cells showing DNA degradation; (**F**) percentage of cells positive for propidium iodide. After 24 h in Gln-free medium, the cells were split into five groups and recovered in the same medium supplemented with different concentrations of Gln (584, 50, 5, and 0.5 µg mL^-1^) or maintained in complete HMI-9 medium (control). At 24 and 48 h of recovery, cells were analyzed again to assess (**G**) polyploidy, (**H**) DNA degradation, and (**I**) membrane integrity. All assays were performed in triplicates (technical replicates), and results represent the mean of three independent biological replicates. Statistical analysis was performed using one-way ANOVA followed by Tukey’s test, with differences considered significant at p < 0.05.

Thus our results show that, regardless of the presence of a GS activity, extracellular Gln at concentrations above 5 μg mL^-1^ is critical for maintaining the cell cycle and thus promoting proliferation. In addition, the deleterious effects of the absence of Gln are reversible within 48 h.

### The absence of Gln affects BSF of *T. brucei*

To further investigate the observed alterations, we used markers of cell cycle (using DNA content as a proxy of ploidy) and cell death (DNA degradation and membrane integrity) to further investigate the role of Gln in cell proliferation. We incubated BSF in Gln-depleted HMI-9 medium, or not (control), for 24 h. For DNA content and DNA degradation analysis, the cells were permeabilized, stained with propidium iodide, and subjected to cytometry analysis. Approximately 22% of the cells maintained in Gln-free medium increased their DNA content, suggesting polyploidy (more than two nuclei per cell) (**Fig. 3 D**). Additionally, the cells did not show DNA degradation when compared to the control (**Fig. 3 E**). When non-permeabilized cells were analyzed for membrane integrity, no differences were observed between the group of cells maintained in Gln-free medium and the control (**Fig. 3 F**). To investigate if the change observed in DNA content in the cells cultivated in Gln-free medium is dependent on the Gln concentration for the recovery of their normal profile, the cells were grown in medium lacking Gln and then transferred to medium supplemented with different concentrations of this amino acid (584, 50, 5 or 0.5 µg mL^-1^). After 24 and 48 h of Gln recovery, the cells were analyzed for plasma membrane integrity, DNA content, and degradation. After 24 h of recovery in different concentrations of Gln, the percentage of cells with higher DNA content decreased from 22% (percentage before resupplying the medium with different concentrations of Gln) to approximately 10%. Only the cells kept at 0.5 µg mL^-1^ Gln maintained above 20% population with the higher DNA content (**Fig. 3 G**). At 48 h, only the cells kept in 0.5 µg mL^-1^ Gln showed around 30% of cells with characteristics of polyploidy, while the cells maintained in 584, 50, or 5 µg mL^-1^ Gln presented a DNA content like the control (**Fig. 3 G**). Regarding markers of cell death, the cells kept in media supplemented with different concentrations of Gln increased the percentage of DNA degradation and loss of membrane integrity after 24 h, when compared to the control (**Fig. 3 H** and **I**). After 48 h, only the cells kept in 0.5 µg mL^-1^ of Gln maintained increased DNA degradation and loss of membrane integrity when compared to the control (**Fig. 3 H** and **I**). These results reinforce our observation that the SMB cells require extracellular Gln at concentrations above 0.5 µg mL^-1^ for the parasite’s survival and correct cell cycle progression.

The observed fact that the absence of Gln triggers an increase in DNA content led us to hypothesize polyploidy (more than two nuclei per cell) and/or polykinetoplasty. Thus, cells incubated in standard or Gln-free media for 24 h were labeled using Hoechst 33342, to stain DNA and analyzed by capturing images using fluorescence microscopy. The number of nuclei and kinetoplasts per cell was counted. The normal K and N configurations in the BSF cells are: 1K1N, 2K1N, and 2K2N (21). Our results show that approximately 50% of the parasites maintained in Gln-free HMI-9 had an abnormal (increased) number of nuclei and kinetoplasts per cell (**Fig. 4**). This data, together with the previous observation about the recovery of normal cell proliferation, confirms that parasites maintained in the Gln-free media were unable to complete the cell division.

**Figure 4.**
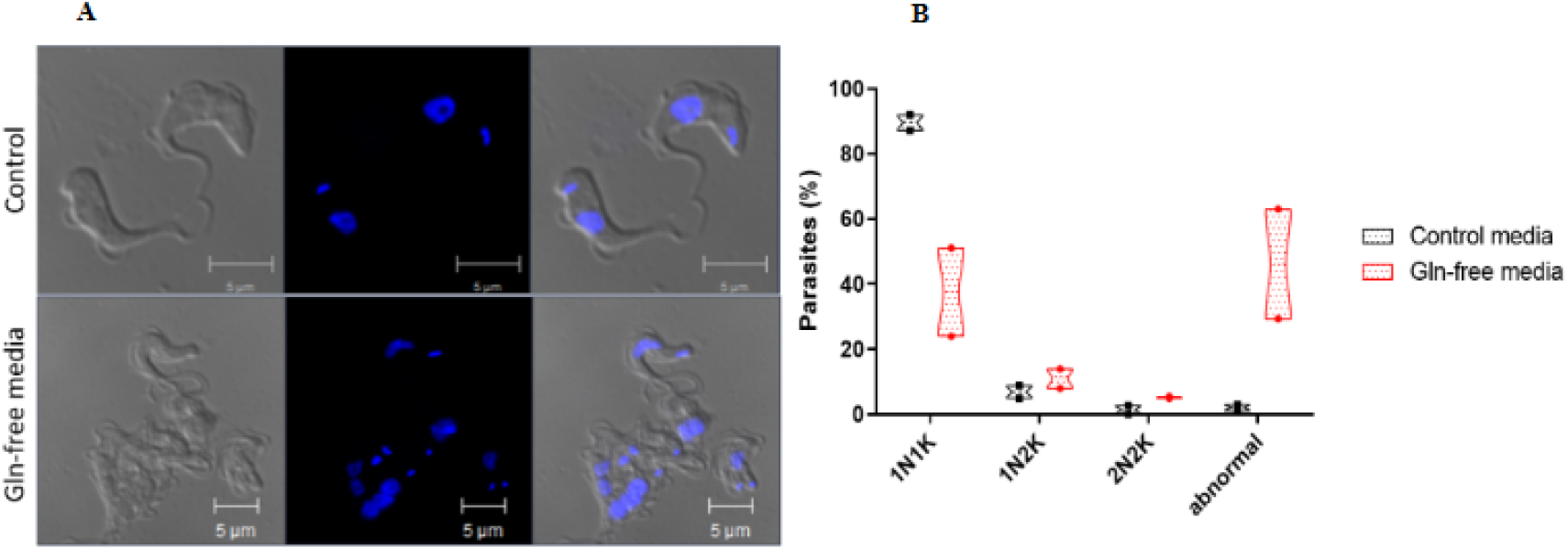
Glutamine deficiency affects the number of nuclei and kinetoplasts in bloodstream forms of *T. brucei*. Cells were maintained for 24 h in HMI-9 media containing or not Gln. (**A)** Stained parasites by Hoechst 33342. (**B)** Classification of the cell population based on the configuration of N and K after different culture conditions.

After detection of abnormal parasites by fluorescence microscopy, we performed an SEM analysis (**Fig. 5 A- D**). The results showed that compared to control cells, cells maintained in the Gln-free HMI-9 presented altered morphology, with a larger cell body and often, more than one flagellum per cell (**Fig. 5 C** and **D**). These data support the idea that the parasites duplicate the nuclei, kinetoplast, and flagellum, but are unable to complete cytokinesis.

**Figure 5.**
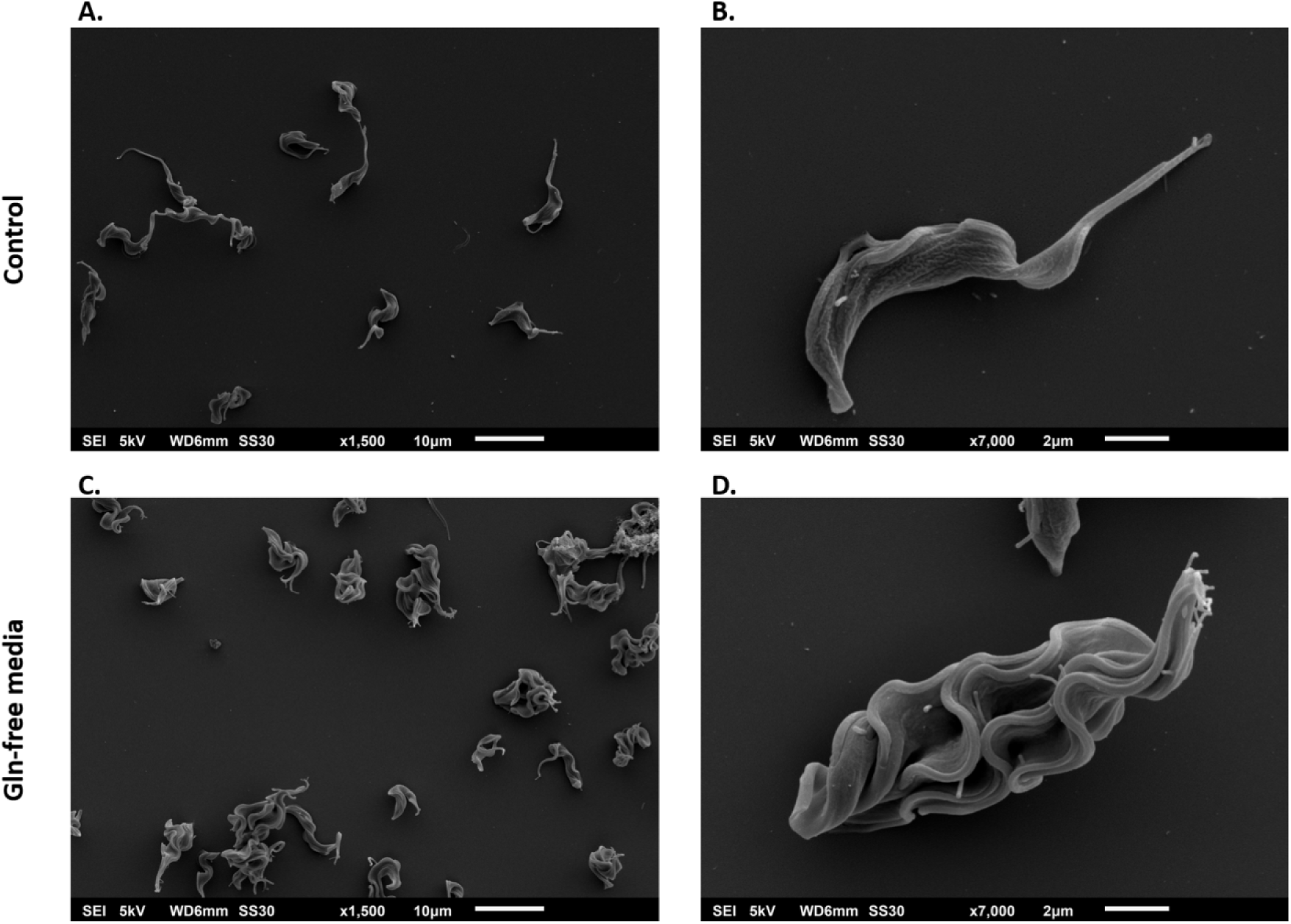
In the absence of Gln, BSF parasites display aberrant cell morphology, as revealed by scanning electron microscopy. BSF of *T. brucei* were maintained in complete HMI-9 media (**Control: A and B**) or without Gln during 24 h (**Gln-free media: C and D**). Then the cells were analyzed by scanning electron microscopy. **A** and **C**: x1,500 magnification of control and tested populations. **B** and **D**: x7,000 magnification of representative parasites of control and tested populations. Size bars: A and C: 10 μm, B and D: 2 μm.

Once confirmed by fluorescence microscopy and SEM that the parasites presented ploidy and morphological alterations when maintained in Gln-free HMI-9, we were interested in analyzing the intracellular ultrastructure of the cells. To assess the intracellular ultrastructure, the cells maintained in the presence or absence of Gln for 24 h were analyzed by TEM (**Fig. 6**). The results confirmed the presence of more than one flagellum per cell, with a single increased nucleus or several nuclei (N) per cell, and many vacuoles, suggestive of lysosomes (**Fig. 6** middle panel). Moreover, when Gln was added back to the culture, the cells rescued the normal morphology and number of structures (**Fig. 6**, bottom panel).

**Figure 6.**
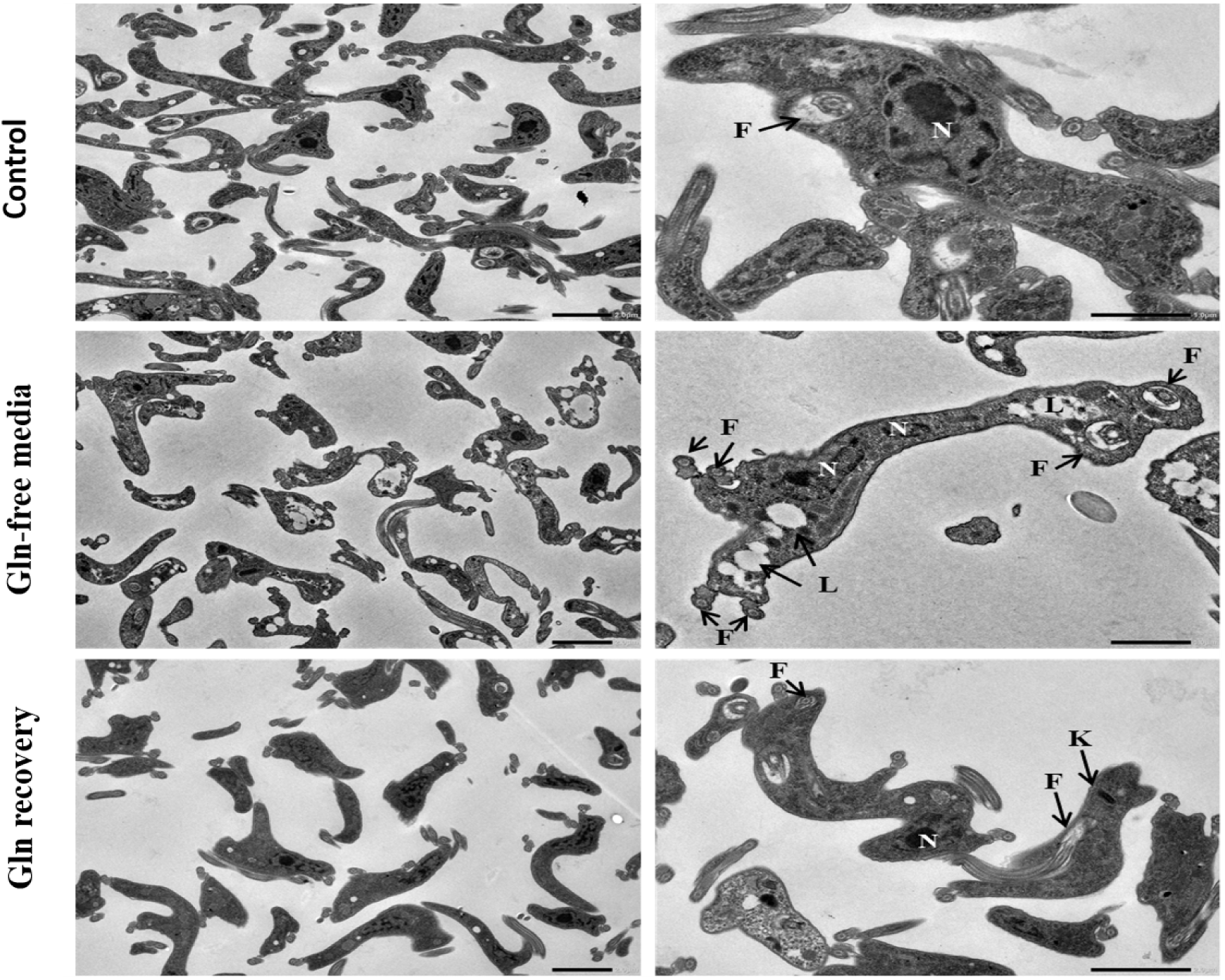
Abnormal cell structures are observed in BSF forms deprived of Gln, according to transmission electron microscopy. BSF of *T. brucei* were maintained for 24 h in Gln-free media or not (control). Gln was added back in the culture, and cells were analyzed 48 h later. **N**: nuclei; **K**: kinetoplast; **F**: flagellum; **L**: lysosome. Left panel: lower magnification, size bars: 2.0 μm (all the pictures). Right panel: higher magnification, size bars: 1.0 μm (upper and middle pictures) and 2.0 μm (bottom picture).

## 3D Cryo-SXT analysis

To further investigate the ultrastructural alterations of cells grown in Gln-depleted or control medium, Cryo-Soft X-Ray Tomography (Cryo-SXT) was employed to visualize in 3D the treated or untreated cells. This approach allows the reconstruction of the 3D structure of the whole intact cell without introducing potential artifacts generated by the physical slicing of the cells (**supplementary material Video A**, **B** and **C**). Our results revealed the presence of multiple organelles and structures that are normally single-copy inside the cells. Of note, multiple nuclear structures and flagella, as well as unusually long and branched mitochondria, were observed (**Fig. 7**). These data are consistent with the observed structures generated as *T. brucei* cells undergo a cell cycle division without completing cytokinesis. Moreover, the statistical information obtained from the Amira 3D software’s Label Analysis tool on the reconstructed volumes enabled us to verify the increase in area and volume of the cells as well as of their organelles (**Fig. 7 and Table 1**).

**Figure 7.**
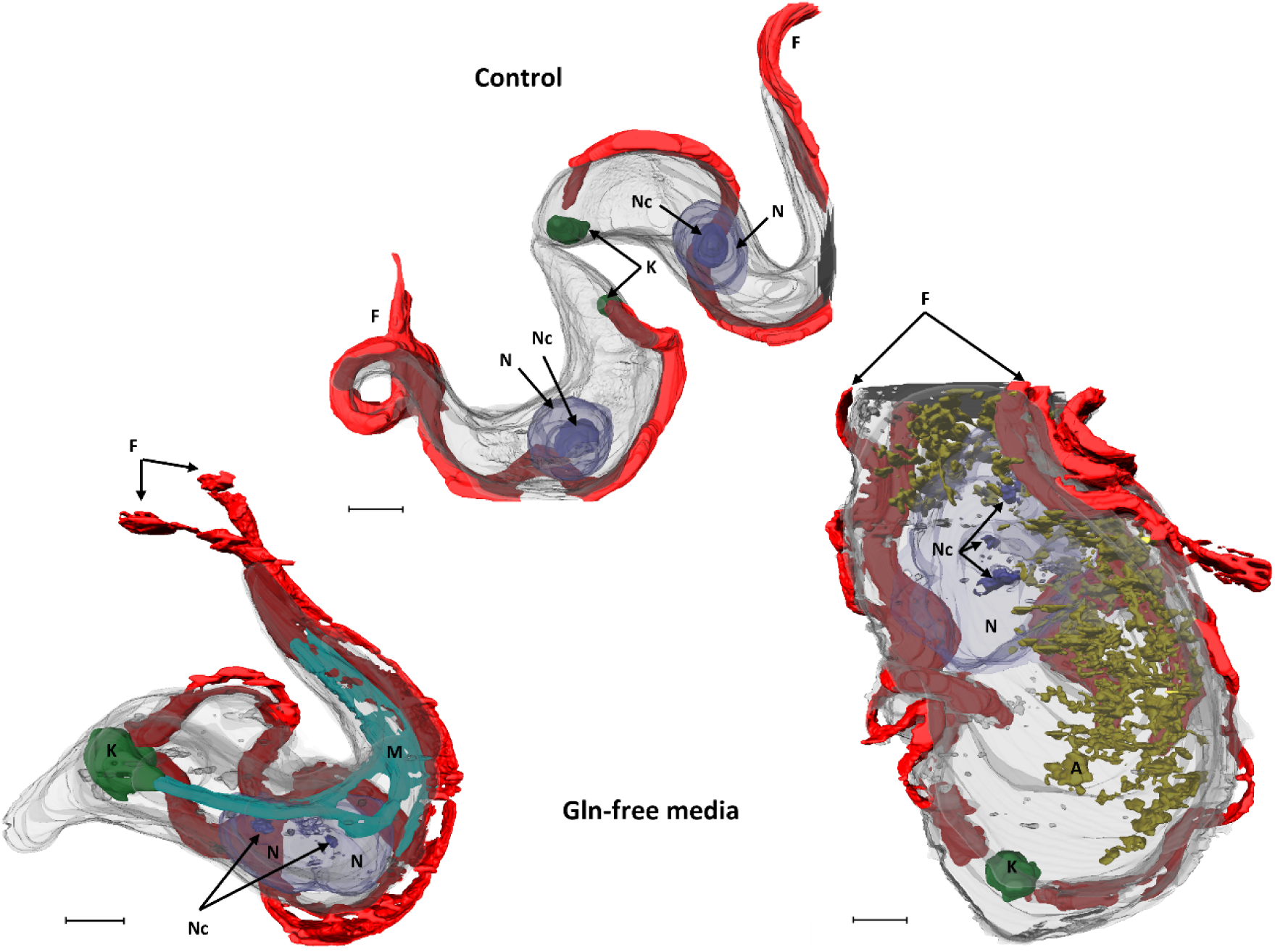
BSF forms deprived of Gln exhibit differences in organellar volume and content, revealed by Cryo-Soft X-ray tomography and reconstruction. BSF of *T. brucei* were cultured for 24 h in Gln-free media or not (control), deposited on Au TEM Quantifoil® grid, and then punge-frozen for Cryo-SXT analysis. The segmentation showed evident differences between both cells, especially considering the increase in number of different organelles as well as changes in size and shape. **N**: nuclei; **Nc**: nucleolus; **K**: kinetoplast; **F**: flagellum; **M**: mitochondria; **A**: dense granules, probably acidocalcisomes. Size bars: 1.0 μm.

**Table 1.** Quantitative analysis of area and volume from BSF of *T. brucei* obtained from segmented Cryo-SXT data by Amira 3D software.

| Cellular feature | Control |  | Gln-free media (left cell) |  |  |  |  | Gln-free media (right cell) |  |  |  |  |
| --- | --- | --- | --- | --- | --- | --- | --- | --- | --- | --- | --- | --- |
|  | Unitary Area (nm<sup>2</sup>) | Unitary Volume (nm<sup>3</sup>) | Unitary Area (nm<sup>2</sup>) | Unitary Volume (nm<sup>3</sup>) | Increase (+) or decrease (-) in size | Change in area (%) | Change in volume (%) | Unitary Area (nm<sup>2</sup>) | Unitary Volume (nm<sup>3</sup>) | Increase (+) or decrease (-) in size | Change in area (%) | Change in volume (%) |
| Cell | 5.04E+03 | 6.04E+03 | 1.24E+04 | 8.30E+03 | + | 145 | 38 | 2.98E+04 | 1.80E+04 | + | 491 | 197 |
| Flagellum | 2.90E+03 | 1.06E+03 | 2.99E+03 | 1.44E+03 | + | 3 | 174 | 2.78E+03 | 1.80E+03 | + | 4 | 71 |
| Nucleus | 7.33E+02 | 6.08E+02 | 5.17E+02 | 4.10E+02 | - | 29 | 33 | 1.61E+03 | 2.88E+03 | - | 119 | 374 |
| Nucleolus | 1.67E+02 | 1.21E+02 | 2.81E+01 | 6.52E+00 | - | 83 | 95 | 6.85E+01 | 1.53E+01 | - | 59 | 87 |
| Kinetoplast | 7.93E+01 | 3.96E+01 | 3.41E+02 | 1.80E+02 | + | 330 | 356 | 1.62E+02 | 8.50E+01 | + | 105 | 115 |

Unitary = means the quantity was normalized by the number of entities present in the cell.

## Glutamine depletion causes a cytokinesis defect

The previous results showed that Gln is relevant for correct cell cycle progression, specifically affecting the G2/M cell cycle phases. To better understand the participation of Gln in the cell cycle, parasites were synchronized in G1 phase. Synchronized cultures were split into two, which were supplemented (control) or not with Gln (**Fig. 8)**. As expected, the cells maintained in the Gln-free medium during 48 h were unable to complete cell division and were arrested as polyploid cells. In contrast, cells maintained in complete medium recovered their normal cell cycle and maintained their proliferative profile (**Fig. 8 A** and **B**). After 48 h of cultivation in the Gln-free medium, the culture was split again into two samples: one was transferred to standard (Gln-supplemented) medium (control), and the other was maintained in the Gln-free medium. The cells incubated in the presence of Gln re-entered the cell cycle in a similar way to controls and were able to properly proliferate after a 72-h recovery (**Fig. 8 A, B,** and **C**). In addition, when the membrane integrity and DNA degradation were assessed as cell death markers, we found that the cells maintained in the Gln-free media lost the membrane integrity and increased the DNA degradation 72 h after cell cycle synchronization, while the cells kept in the complete medium maintained a normal profile (**Fig. 8 D** and **E**). The results show that the synchronized cells maintained in the Gln-free medium were able to go from G1 to S, G2/M phases and polyploid cells, but could not complete the cell division, showing that Gln is relevant to cytokinesis. Summarizing, the parasites duplicated the genetic material and reentered the cell cycle but could not complete cell division, thus becoming polyploid. Interestingly, when Gln was added back to the media, the cells recovered the normal proliferative profile and the correct cell cycle progress. These data reinforced the finding that, although the cells can biosynthesize Gln, this is not enough to supply the parasite’s Gln requirements.

**Figure 8.**
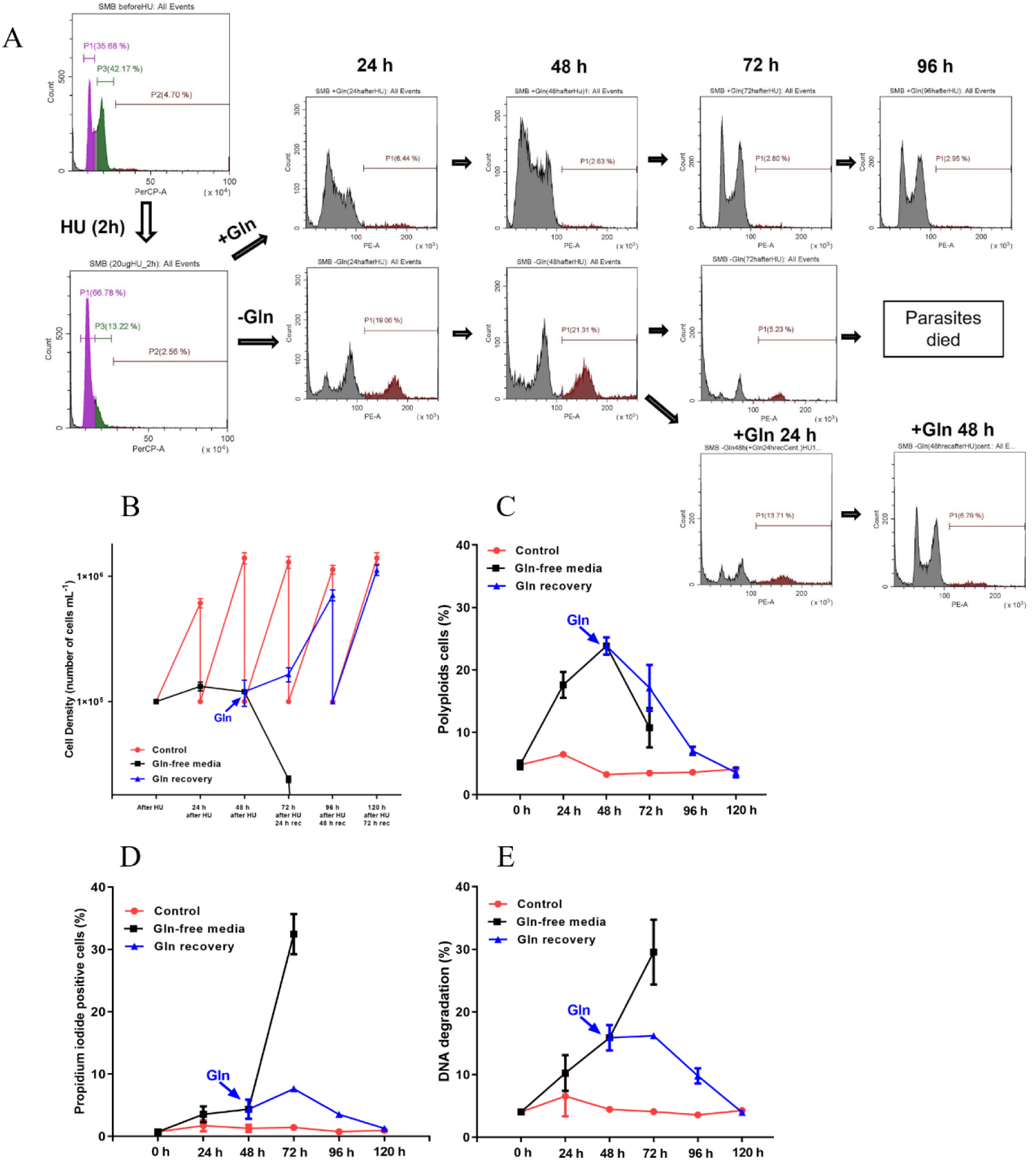
Cell cycle defects in BSF forms lacking Gln are demonstrated after hydroxyurea treatment and flow cytometry analysis. **(A)** Histograms of the cell cycle analysis: the cells were treated with hydroxyurea, after treatment, the cells were split into two cultures with (+Gln) or without (-Gln) Gln during 48 h. Then, the culture was split into two cultures again, and Gln was added back in the media. **(B)** Cell density: every day the cells were counted and diluted back to 1 x 10^5^ mL^-1^. **(C)** Quantification of polyploid cells present in the cultures every day, **(D)** Quantification of cells that lose the membrane integrity, **(E)** DNA degradation. These parameters were analyzed by flow cytometry each 24 h. (**Red line**: control; **Black line**: Gln-free media; **Blue line**: Gln recovery).

## Absence of glutamine impairs cell motility

The absence of Gln affects cell morphology, generating aberrant multiflagellated parasites with an enlarged cell body. It is conceivable to hypothesize that those cells have decreased motility when compared to controls. To measure cell motility, the parasites were maintained in standard medium (as a control) and Gln-free medium for 24 hours. Then, cell motility was monitored in a spectrophotometer. A reduction of the absorbance at 600 nm due to sedimentation (sedimentation assay) is inversely related to the cell motility, if it is assumed that the ability of cells to remain in suspension reflects the motility of the flagellum. Parasites maintained in Gln-free medium were unable to swim in the culture, progressively clustering at the bottom of the cuvette. This sedimentation defect is likely associated with impaired flagellar motility or could be related to the increase in cell density, as demonstrated in previous assays. In contrast, control cells remained homogeneously distributed in the medium, as expected for normal motility (**Fig. 9**). Altogether, these results indicate that in the absence of Gln, BSF parasites exhibit reduced motility and adopt a tumble-cell phenotype.

**Figure 9.**
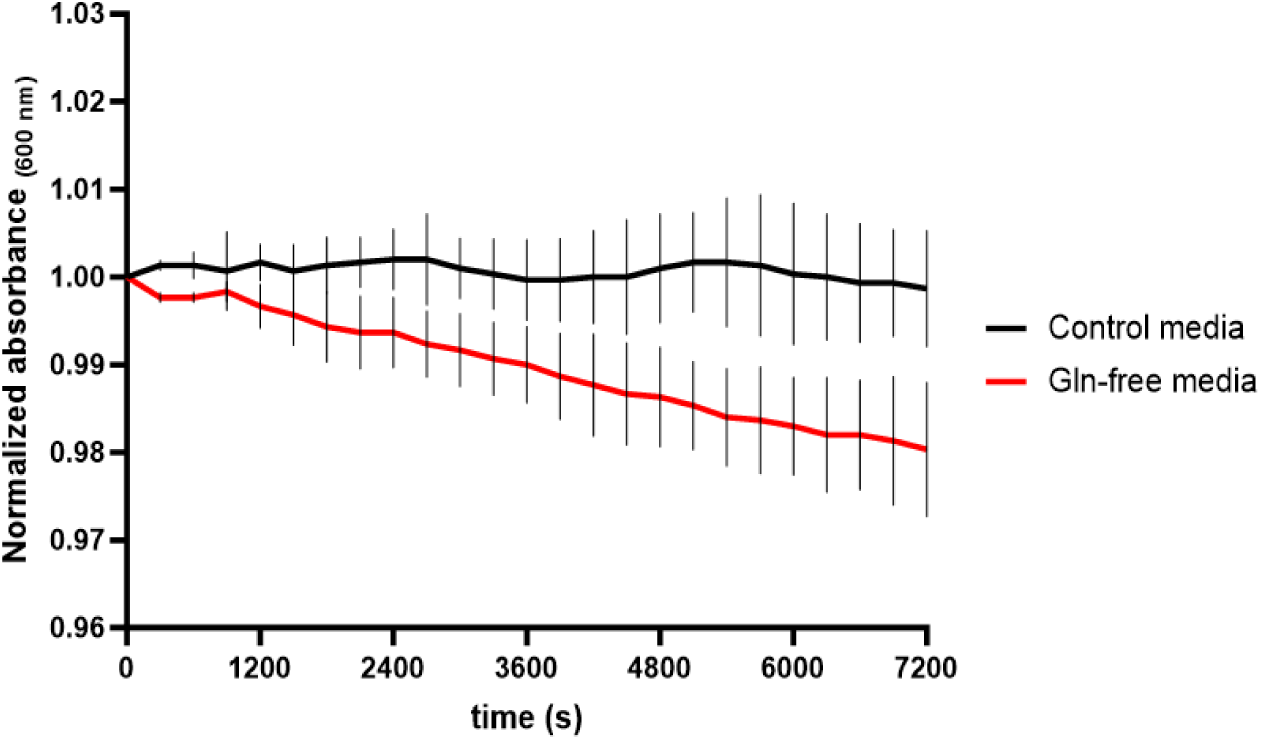
**In the absence of Gln, BSF parasites show decreased motility**. The motility of BSF cells was measured using a sedimentation assay. The optical densities (OD600) of the cell suspension maintained in standard medium or Gln-free medium for 24 h were monitored by spectrophotometry. **Black line**: control; **Red line**: without Gln for 24 h.

## Supply of glutamine is important to protein glutamylation

Glutamylation and polyglutamylation are important PTMs to regulate cytoskeletal dynamics. In decreased glutamylation levels, PCF of *T. brucei* exhibit defects in growth and cytokinesis as well as impaired cell motility (14,15). Given the morphological and functional changes detected in parasites grown in the absence of Gln, observed here and the relevance of these PTMs to parasite development as documented in the literature (12,14,15), we investigated the participation of Gln in the glutamylation process in the *T. brucei* BSF. For this purpose, the cells were maintained in medium with or without Gln for 24 h. To analyze the recovery from Gln deprivation on protein glutamylation, samples of cells maintained in Gln-free medium were transferred to medium containing Gln for 48 h. The glutamylation and/or polyglutamylation levels of the cells were initially analyzed by western blot and immunofluorescence using the antibodies GT335 and PolyE. The immunofluorescence analysis revealed glutamylation and/or polyglutamylation of proteins when the parasites were kept in standard culture conditions (**Fig. 10 A**, control panel). After 24 h without Gln, it was possible to observe a decrease in the fluorescence signal from both GT335 and PolyE markers, indicating that these parasites have lower levels of glutamylation (**Fig. 10 A**, Gln-free medium). When Gln is added back to the medium, the levels of glutamylation are restored, indicating that the external supply of Gln is crucial for proper glutamylation in *T. brucei* (**Fig. 10 A**, Gln recovery). To corroborate these observations, we detected the levels of glutamylation by western blot using total protein extracts. As previously indicated, the parasites maintained in the Gln-free medium showed lower levels of glutamylation (**Fig. 10 B**). Our data showed for the first time that the purvey of exogenous Gln effectively participates in the glutamylation process in *T. brucei* BSF forms.

**Figure 10.**
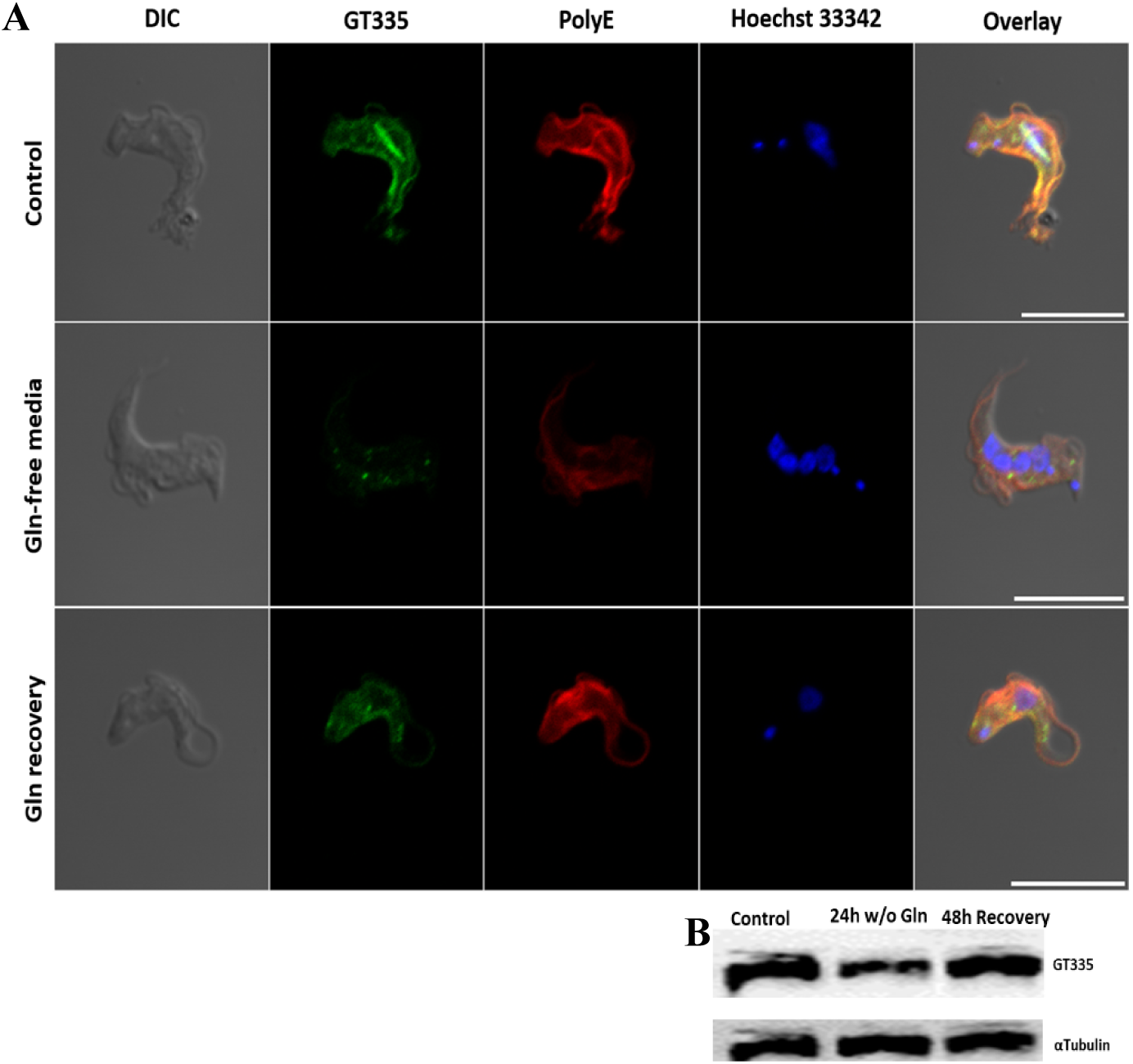
BSF parasites cultivated in the absence of Gln show decreased levels of glutamylation. Bloodstream forms of *T. brucei* were maintained in HMI-9 complete medium (control) or in the presence of Gln-free medium for 24h. After 24h of cultivation, Gln was added back into the media, and the cells were maintained for 48h. The glutamylation and polyglutamylation levels of the cells were analyzed using the antibodies GT335 and anti-polyglutamate chain (polyE), and the cell nucleus was stained using Hoechst 33342. (**A**) Immunofluorescence analysis; (**B**) Western blot analysis using the antibody GT335.

Our results show that exogenous Gln is fundamental for protein glutamylation. As it has been reported that the subpellicular and flagellar microtubules are among the most glutamylated proteins in *T. brucei* PCF (16), we considered it relevant to analyze the glutamylation of the cytoskeleton and its associated proteins in BSF in more detail. For this, an enriched cytoskeleton fraction of the parasites was obtained and analyzed by mass spectrometry. A total of 65 glutamylated proteins were detected, 32 of them are annotated as hypothetical proteins, and 33 have attributed functions in their annotation (**Fig. 11 A**; **supplementary material Table 1**). The identified glutamylated proteins in the cytoskeleton-enriched fraction were glutamylated in one or more positions: 35 out of 65 (53.8%) had 1 glutamylation site and 14 out of 65 (21.53%) had 2 glutamylation sites (**Fig. 11 B**). Among the identified proteins we found, alpha-tubulin and many other proteins involved in different biological processes. Specifically, it was possible for alpha-tubulin to detect glutamylation at residue E445, as previously described for PCF (12,16). In our experiments, we could also observe that other glutamate residues are glutamylated in the protein, such as E438, E446, E449, and E450. When compared to the alpha-tubulin from *Chlamydomonas reinhardtii* (95% similarity in amino acid sequence), it was possible to observe that some glutamylation sites are conserved (E445, E449, and E450). However, *T. brucei* alpha-tubulin showed two previously undescribed glutamylation sites (E438 and E449) (**Fig. 11 C**). The Gene Ontology (GO) enrichment analysis from the TriTrypDB database linked glutamylated proteins in the cytoskeleton fraction to several biological processes. These proteins are associated with microtubule organization, the mitotic cell cycle, anaphase, cell motility, intraciliary transport, and defense against host immune responses. They also play a role in the metabolism of galactose, glucose-6-phosphate, oxaloacetate, NADH, NADPH, malate, carbohydrates, and nucleotide sugars (**Fig. 11 D**; see complete list in the **supplementary material Table 2**). Considering that the analysis was made on a cytoskeleton fraction, the most abundant glutamylated proteins detected were part of microtubules, axonemes, and other cytoskeleton components. These results are consistent with our previous results, revealing a relationship between Gln scarcity in the culture medium and defects in cell division and motility. The GO enrichment molecular function analysis suggested that glutamylation might be important for proteins with different functions within the cell, such as the regulation of the GTPase, UDP-glucose 4-epimerase, L-malate dehydrogenase, lipid transport activity, tRNA-uridine aminocarboxypropyltransferase activity. Also, glutamylation might be critical for other essential cellular activities, including ribonucleotide and nucleotide binding; purine ribonucleoside (**Fig. 11 E**; **supplementary material Table 3**). Based on GO enrichment cellular component analysis, we found a widespread distribution of glutamylated proteins in the cell beyond microtubules, including glycosomes, peroxisomes, ciliary pocket, plasma membrane, and the mitochondrial endopeptidase Clp complex (**Fig. 11 E; supplementary material Table 4**). Overall, our data show for the first time that glutamylation is a PTM found in a broad range of proteins in BSF of *T. brucei*.

**Figure 11.**
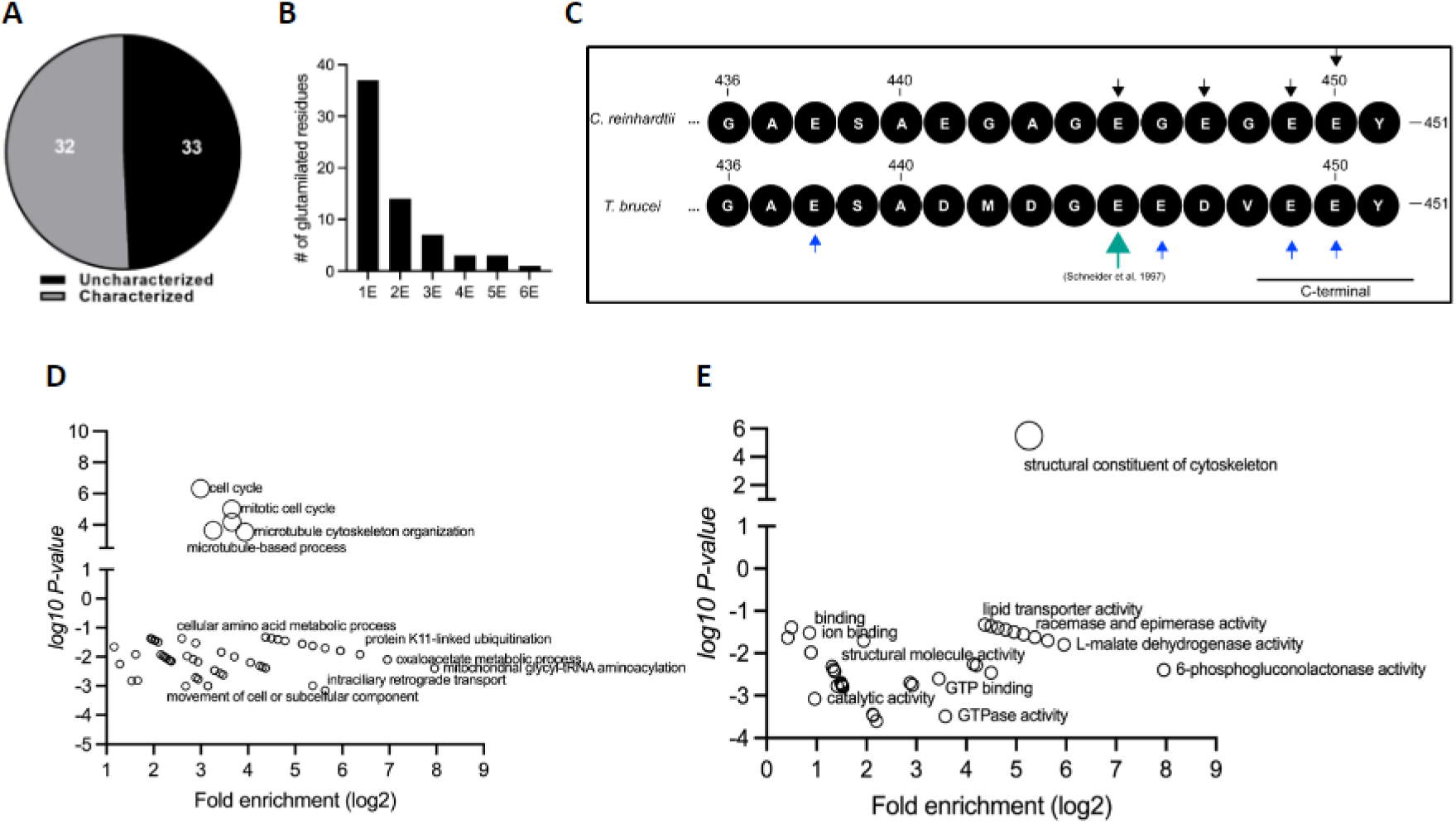
Proteomic profiling of glutamylated proteins in cytoskeleton-enriched fractions of *T. brucei* BSF. BSF of *T. brucei* were maintained in HMI-9 medium and the enriched cytoskeleton fraction was extracted and analyzed by mass spectrometry. **(A)** Pie chart indicating the amount of glutamylated proteins detected. **(B)** Number of glutamylated residues in the glutamylated proteins detected. **(C)** Representation of glutamylation sites (arrows) detected in alpha-tubulin enriched extracts of BSF of T. brucei and comparison with *C. reinhardtii*. **(D)** GO enrichment biological function. **(E)** GO enrichment molecular function. To check the complete list of glutamylated proteins, see the supplementary material.

## Discussion

In mammals, the BSF of *T. brucei* can survive in blood, central nervous system, and interstitial spaces of adipose tissue, among other territories (22). In blood, Gln is the most abundant free amino acid, with a concentration of approximately 700 µM. In contrast, the concentration of its precursor Glu is approximately 20 µM (23). Therefore, as an extracellular parasite, *T. brucei* is always exposed to the Gln present in the bloodstream. Gln can feed multiple pathways in the BSF of *T. brucei*, including several involved in mitochondrial metabolism. Additionally, it constitutes the primary source of amino groups for the nitrogen pool of the parasite (11). Here, we describe unexpected roles for Gln in cell cycle regulation and its participation as a substrate for protein glutamylation in BSF of *T. brucei*. Our results show that BSF can obtain Gln from the external medium and through its biosynthesis from Glu using a GS. However, the partial depletion of GS by RNAi did not impair the ability of BSF to undergo normal cell proliferation when maintained in Gln-rich medium. This is in agreement with previous observations on the essentiality of Gln and on the role of this amino acid in the biosynthesis of Glu (11). Importantly, BSF maintained in Gln-depleted medium were unable to complete cytokinesis, resulting in the generation of aberrant polyploid cells due to cell cycle arrest and the absence of cell division progression. As occur with most of eukaryotic cells, the cell cycle in *T. brucei* goes throughout the classical G1, S, and G2/M phases (24) with some unique characteristics: while canonical Cdc6 and Cdt1 regulators are absent, some others (which have been considered as trypanosomatid-specific) are present in the parasite, such as cyclins, cyclin-dependent kinases, and mitotic centromere-associated kinesins (25). It has been established that the cell cycle starts with the elongation and maturation of the parabasal body, with the initiation of the formation of a new flagellum. Next, the replication of kDNA and gDNA starts, initiating nuclear and mitochondrial replication (24,25). In this context, proper flagellar motility is essential for cytokinesis (26). A successful cell division ends with cells having one nucleus and one kinetoplast (K1N1) each. Cells having two kinetoplasts and one nucleus (K2N1) correspond to cells in late G1/early S, while cells having two kinetoplasts and two nuclei (K2N2) are in late S or early M phase (21). Our results show that cells maintained in low Gln medium showed an increased number of polyploid cells with an abnormal number of K and N. Additionally, our results show that cells in the G1 phase, when maintained in low Gln medium, undergo S and G2/M phases but are unable to complete cell segregation, explaining the finding of polyploid parasites. Importantly, even undergoing these structural alterations, the cells maintained for 48 h in Gln-depleted medium kept their membrane and DNA integrity. When Gln was added back to the culture, they recovered their regular proliferation and structure.

It is known that BSF of *T. brucei* can re-enter a new cell cycle before completing cytokinesis, suggesting that the cells lack a checkpoint between cytokinesis and the new G1 phase, which results in deficient cellular segregation and polyploidy (27). Our results, obtained using SEM and TEM, showed that BSF maintained in Gln-depleted medium became multi-flagellated, lacked the ingression of a cleavage furrow, and became larger than untreated cells. The ingression of a cleavage furrow is the last step to cell segregation, and it occurs when the organelles (nuclei, kinetoplast, and flagella) are completely duplicated prior to the cell segregation into two daughter cells. This process depends on the remodeling of microtubules, the cytoskeleton, and cell membranes (24). The fact that when Gln was added back, the parasites rescued normal cell cycle progression and proliferative profile suggests that this amino acid plays a crucial role in cell cycle coordination.

In trypanosomatids, the cytoskeleton is composed only of microtubules. These microtubules are responsible for the cell shape and constitute the mitotic spindle, the flagellar axoneme, the basal body of the flagellum, and the subpellicular corset (28). Additionally, the cytoskeleton plays a crucial role in various cellular processes, including the positioning of organelles, intracellular transport, mitosis, cytokinesis, and cell motility (14,18,19). The diversity of microtubules is responsible for the large variety of their cellular functions, and is generated in part by PTMs, such as acetylation, phosphorylation, polyglutamylation, polyglycylation, palmitoylation, polyamination, and detyrosination (13,17). Among these PTMs, polyglutamylation and glutamylation are described as important to keep the microtubules and organelles in the correct position in eukaryotic cells, including trypanosomatids (14,29–31). Our results showed that BSF cells maintained in Gln-depleted medium reversibly decreased the levels of glutamylation and polyglutamylation of microtubules. The levels of glutamylation returned to normal when Gln was added back into the medium. These results indicate that Gln actively participates in the glutamylation and polyglutamylation process in BSF of *T. brucei*. Our findings, along with recent reports (11), show that in BSF *T. brucei*, Glu for protein glutamylation largely derives from extracellular Gln.

Glutamylation and polyglutamylation are described as adding one or more Glu residues within the primary sequence of the target protein, essentially alpha- and beta-tubulin, generating side chains of variable length (32). Although the family of tubulin tyrosine ligase-like (TTLL) enzymes is described as responsible for adding Glu residues in tubulin, the biochemical characteristics of these enzymes are still unknown (12,16,17,33,34). Previous studies have demonstrated the importance of glutamylation to procyclic forms of *T. brucei*: flagellar microtubules and subpellicular microtubules form a corset that is substantially glutamylated, and almost the entire cytoskeleton is glutamylated (16). Knocking down the enzyme TTLL4B (a homolog of a TTL protein in *T. brucei*) induced a decrease in cell growth rate, with blockage of cytokinesis and decreased tubulin glutamylation levels (12). Recently, it was demonstrated that polyglutamylation regulates cytoskeletal architecture and motility of the procyclic forms.

The depletion of polyglutamylases TTLL6A or TTLL12B perturbs cytoskeletal architecture, leading to diffuse motility, concomitant with a reduction of axonemal polyglutamylation (14). TTLL1 is essential for the integrity of microtubule structure and polyglutamylation but does not affect cellular motility (15). The role of glutamylation in regulating axonemal structure, motility, ciliary transport, and extracellular vesicles has been described in other organisms, such as *Chlamydomonas reinhardtii* (35,36) and *Caenorhabditis elegans* (31). In HeLa cells tubulin polyglutamylases are regulated during the cell cycle, presenting polyglutamylase activity peaks in G2, while tubulin glutamylation is highest in mitosis (37). Although tubulin is the most described glutamylation target, other proteins can also be glutamylated. In a similar fashion, a broad range of proteins, such as nucleoporins and nucleosome assembly proteins (NAP1 and NAP2), are substrates for glutamylation in HeLa cells (38).

Our mass spectrometry analysis of enriched cytoskeleton fractions demonstrated a broad variety of glutamylated proteins in BSF, including, as expected, tubulin. Remarkably, the group of glutamylated proteins includes proteins involved in cellular regulation as pantothenate kinase, the first enzyme of the coenzyme A biosynthetic pathway, and AMPK1, responsible for the balance between proliferation and differentiation in BSF (39), phosphatidylinositol 3-kinase, an enzyme required for normal proliferation and endocytosis in *T. brucei* (40), and a variant surface glycoprotein (VSG), important for the parasite to evade the host immune system. Another group of glutamylated proteins includes the DNA polymerase and RNA pseudouridylate synthase. Furthermore, proteins involved in metabolism, such as malate dehydrogenase (TCA cycle), phosphogluconolactonase (pentose phosphate pathway), UDP-glucose 4-epimerase, and ABC transporters, are also glutamylated. In summary, our data demonstrate that in BSF forms of *T. brucei*, a variety of proteins are also substrates for glutamylation, indicating their possible participation in multiple processes such as microtubule dynamics, gene expression and regulation, cell cycle, and metabolism. Further studies are required to understand the role of glutamylation and polyglutamylation in BSF forms of *T. brucei*. It is important to note that the mass spectrometry analysis was performed using an enriched cytoskeleton fraction, which explains most of the identified glutamylated proteins being related to cytoskeleton structure. Possibly, using total cell extracts would reveal additional glutamylated proteins in the future. Further analyses need to be applied to identify the number of glutamylated proteins and the function of glutamylation and polyglutamylation in the BSF of *T. brucei*.

Until now, Gln has not been described as a player in the glutamylation process. All data in the literature describe Glu as the amino acid involved in this PTM, both in *T. brucei* PCF and other eukaryotic cells (12,16,29). To our knowledge, our studies are the first demonstrating the importance of Gln for glutamylation processes and cell cycle regulation in *T. brucei* BSF. It is essential to note that Glu was always present in the cultivation media. As the FBS was treated with commercial glutaminase, Gln was the only amino acid depleted from HMI-9 medium. This fact reinforces that the biosynthesis of Gln from Glu, if operative, is insufficient to supply the parasite with the substrate necessary for energy metabolism and PTMs. Thus, the *T. brucei* BSF parasite relies solely on the external supply of Gln in the medium.

In conclusion, our data show that Gln uptake is essential for proliferation, cell cycle progression, and glutamylation in BSF of *T. brucei*. Glutamylation and polyglutamylation seem to be important for microtubule dynamics, which could explain the impact of glutamine scarcity on the cell cycle in *T. brucei* BSF. Given that Gln is the most abundant amino acid in human blood and that the *T. brucei* BSF lives in the bloodstream, we suggest that the Gln uptake could be a promising target for the development of new therapies against African trypanosomiasis.

## Materials and methods

### Parasites and cell culture

BSF of *T. brucei brucei* (Lister 427) were maintained with successive passages in HMI-9 media, 10% FBS, at 37 °C in a water-saturated atmosphere in the presence of 5% CO_2_.

### Depletion of Glutamine in the Fetal Bovine Serum

To deplete the Gln content in the FBS, the serum was treated with glutaminase (Sigma-Aldrich) following the manufacturer’s instructions. After treatment, depletion of Gln was confirmed by liquid chromatography (*LC* 3000 Eppendorf-Biotronik equipment). The Gln-depleted serum was used to supplement the Gln-free HMI-9 media.

### Knockdown of the glutamine synthetase in BSF of *T. brucei brucei*

#### Transfection of parasites

The glutamine synthetase (GS) putative gene (Tb927.7.4970) was cloned into the tetracycline-inducible vector p2T7-177, using phleomycin as a marker of selection in the culture medium. The construction was confirmed by restriction enzyme analysis. The correct gene sequence was confirmed by PCR using specific primers and sequencing. After confirmation, BSF in exponential growth phase (1.5 x 10^6^ mL^-1^) were transfected with the construction p2T7*Tb*GS. The transfected parasites were selected by the addition of phleomycin into the culture. After the selection of parasites, the *Tb*GS knockdown was induced by adding tetracycline (2 µg mL^-1^) to the culture medium. BSF cells were cultivated in HMI-9 medium at an initial concentration of 1 x 10^4^ mL^-1^.

#### qRT-PCR to confirm knockdown

After 48 h of induction, the RNA was purified using the RNeasy kit from Qiagen, and the cDNA was synthesized using the RevertAid H minus kit from Thermo Scientific, according to the manufacturer’s instructions. The cDNA was applied for qRT-PCR with *Tb*GS-specific primers to analyze the levels of its transcripts. Parasites without transfection and uninduced BSF p2T7*Tb*GS were used as controls.

## Glutamine and glutamate transport assay

BSF in the exponential growth phase were used to evaluate the uptake of Gln and Glu from the external media, as described by (41). Briefly, 1 × 10^7^ parasites diluted in 100 µl TDB (*Trypanosoma Dilution Buffer*: 5 mM KCl, 80 mM NaCl, 1 mM MgSO_4_, 20 mM NaH_2_PO_4_, 20 mM D-Glucose) were placed in 1.5 mL tubes. Uptake studies were initiated by the addition of 3 mM radiolabeled Gln (L-[3,4-3H(N)]-Gln) or Glu (L-[3,4-^3^H]-Glu) (Perkin-Elmer, Boston, MA, USA) and incubated for different times, at 37 °C. Transport was stopped by the addition of 800 µl of ice-cold stop solution (50 mM Gln diluted in PBS). The parasites were washed twice in PBS, resuspended in scintillation liquid (Optiphase Hisafe III, Perkin-Elmer), and analyzed in a scintillation counter (Perkin-Elmer Tri-Carb 2910 TR). The incorporated Gln was calculated as follows: 𝐺𝑙𝑛_𝑖_ = 𝑐𝑝𝑚_𝑖_ [𝐺𝑙𝑛] ʋ cpm^―1^where Gln_i_ is the intracellular Gln; cpm_i_ is the number of cpm associated with the cells due to the uptake of the radiolabeled substrate; [Gln] is the nanomolar concentration of radiolabelled Gln; ʋ is the volume of radiolabelled Gln; and cpm_st_ is the total cpm measured for the added radiolabelled Gln. The graphs, statistical analyses, and data adjustments were performed using the program GraphPad Prism (5.01), GraphPad Software, San Diego, California, USA.

## Glutamine synthetase activity assay

The GS activity was measured in total protein extracts of BSF following the protocol described by Crispim and colleagues (20). In brief, parasites in the exponential proliferation phase were washed twice in PBS and resuspended in lysis buffer (50 mM Tris; 0.25 M sucrose; 0.1 M NaCl; 0.2% triton X-100; pH 7.6) in the presence of 1% protease inhibitors (E-64, TLCK and PMSF) and lysed by freezing (liquid nitrogen) and thawing at 37 °C (5 cycles). Then lysed cells were centrifuged (30 min, 7,000 *x g* at 4 °C) and the supernatants were used to measure GS activity. As the GS reaction consumes ATP (20), we calculated the produced ADP by applying a coupled enzyme assay, as shown (Scheme 1). Here, oxidation of NADH, equivalent to the concentration of consumed ATP, was monitored by measuring absorbance at 340 nm using a spectrophotometer (Evolution 300, Thermo Scientific). Various amounts of the soluble fraction of total protein extracts of BSF (10, 25, and 50 µg) were used. The NADH consumption was quantified based on its molar extinction coefficient (ε = 6,200 M⁻¹ cm⁻¹).

**Scheme 1.**
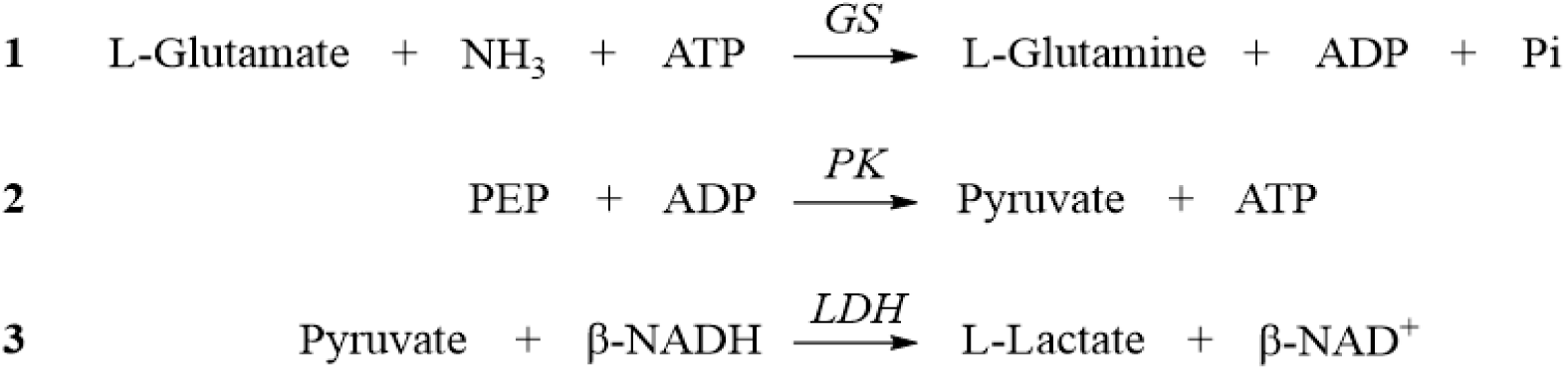
**Schematic representation of the coupled enzymatic reactions used for glutamine synthetase activity assay**. The reaction sequence includes: (**1**) Glutamine Synthetase (GS), which catalyzes the ATP-dependent conversion of L-glutamate and ammonia into L-glutamine; (**2**) Pyruvate Kinase (PK), which regenerates ATP by transferring a phosphate group from phosphoenolpyruvate (PEP) to ADP, producing pyruvate; and (**3**) Lactate Dehydrogenase (LDH), which reduces pyruvate to L-lactate while oxidizing β-NADH to β-NAD⁺.

### Effect of a gradual decrease of the glutamine concentration on bloodstream forms proliferation, cell cycle progression, and membrane integrity

To evaluate the importance of Gln for the parasités survival, BSF were grown in HMI-9 (initial concentration: 1 x 10^5^ cells mL^-1^) with daily dilution to 1x10^5^ cells mL^-1^ again and a successive decrease of the Gln concentration in the media each day (50, 5, 0.5 µg mL^-1^). After cultivation in low glutamine (Gln) medium (0.5 µg mL^-1)^ for three days, the parasites were transferred to complete HMI-9 containing the standard Gln concentration (584 µg mL^-1^) and cultivated for an additional 48 hours. Cell density was determined daily by counting BSF in the Neubauer chamber; aliquots were then separated to assess cell-membrane integrity, cell cycle progression, and DNA degradation, as described in the following sections.

### Cell cycle and DNA degradation

To assess the DNA content (to calculate cell cycle progression and DNA degradation), 1 x 10⁶ cells were washed once with PBS, centrifuged (1,000 x *g*, 5 min), resuspended in 100 µL lysis buffer (64 µM digitonin in phosphate buffer: 7.7 mM Na_2_HPO_4_; 2.3 mM KH_2_PO_4_; pH 7.4), and incubated on ice for 30 min. After incubation, 100 µL of propidium iodide solution (20 µg mL^-1^) was added, and the lysates (5,000 events per sample) were analyzed by flow cytometry, using the channels PE-A and Count (CitoFLEX, Beckman Coulter). The cell cycle phases were analyzed by measuring the DNA content corresponding to G1, S, and G2/M. A DNA content corresponding to sub-G1 was considered as DNA degradation, and a DNA content higher than G2/M was considered as polyploidy. The measurements were performed in triplicate, and data corresponding to the mean were obtained by measuring three independent biological replicates. Statistical analysis was made by performing a one-way ANOVA followed by Tukey’s test. P < 0.05.

### Membrane integrity

To assess the cell-membrane integrity, parasites were collected by centrifugation (1,000 x *g*, 5 min, RT) and washed once in annexin-binding buffer (10 mM HEPES; 140 mM NaCl; 5 mM CaCl_2_, pH 7.4). The parasites were resuspended in 50 µL of the same buffer supplemented with 1 µg µL^-1^ propidium iodide, incubated on ice for 15 minutes, and diluted 10X by addition of 450 µL of annexin buffer. The cells were analyzed by flow cytometry (5,000 events per sample) in a CitoFLEX, Beckman Coulter cytometer using channel PE-A and Count. Positive cells were defined as those that lost membrane integrity. As a positive control for loss of membrane integrity, parasites were treated with digitonin. The measurements were performed in triplicate, and data corresponding to the mean were obtained by averaging the results from three independent biological replicates. Statistical analysis was made by performing a one-way ANOVA followed by Tukey’s test. P < 0.05.

### Recovery at different concentrations of Glutamine

To assess if the recovery of parasites depends on the Gln concentration, BSF cells were incubated in Gln-free medium for 24 h, before the parasites were transferred to HMI-9 media supplemented with different concentrations of Gln (584, 50, 5, 0.5 µg mL^-1^) for 48 h. Cell cycle progression, DNA degradation, and membrane integrity were analyzed daily by flow cytometry. The number of parasites was determined daily by Neubauer chamber count, before cell density was adjusted to 1 x 10^5^ cells mL^-1^ again.

### Cell cycle synchronization

To synchronize the parasite’s cell cycle, exponentially proliferating BSF were treated with hydroxyurea (20 µg mL^-1^) (42) for 2 h. After incubation, the synchronization was confirmed by flow cytometry analysis, and the hydroxyurea was removed by centrifugation. Then, synchronized parasites (1 x 10^5^ mL^-1^) were resuspended in HMI-9 Gln-free medium or complete medium as a control. After 48 h of cultivation in the Gln-free HMI-9 medium, the culture was split into two flasks; one was transferred to complete HMI-9 medium, while the other was maintained in the Gln-free medium. The culture was followed up for 5 days, analyzing cell cycle, membrane integrity, and number of cells every day, before adjusting cell density to 1 x 10^5^ mL^-1^.

### Western blot analysis

To analyze the participation of Gln in glutamylation of tubulin (PTM), the parasites (1 x 10^7^) were maintained in HMI-9 (complete or Gln-free) for 24 h. After 24h in the Gln-free medium, Gln (584 µg mL^-1^) was added back to the culture for 48 h. After respective incubations, the parasites (1 x 10^7^) were resuspended in lysis buffer (phosphate buffer + protease inhibitors) and lysed by freezing (liquid nitrogen) and thawing at 37 °C (5 cycles). After lysis, the parasite extracts were resuspended in SDS sample buffer (62.5 mM Tris/HCL, pH 6.8; 2,3% SDS; 10% glycerol; 0.01% bromophenol blue; 20 mM mercaptoethanol) in the proportion 4/1, then the samples were boiled for 5 min at 100 °C. The total protein extracts were analyzed by SDS-PAGE and transferred to the nitrocellulose membrane. The membrane was blocked with 5% of free-fat milk solution (diluted in PBS + 0.01% Tween20) and incubated with a 1:5,000 dilution of the mouse antibody GT335 (recognizing polyglutamylated proteins, from AdipoGen^®^) prepared in 3% of free-fat milk solution for 1 h at room temperature. After incubation, the membrane was washed and incubated with a 1:10,000 dilution of goat antibody anti-mouse IgG-HRP conjugate (Bio-Rad) prepared in 3% fat-free milk solution for 1 h at room temperature (RT). Then the membrane was washed and incubated with Pierce™ ECL Western Blotting Substrate (Thermo Scientific) for 5 min. The signal was detected using the equipment ChemiDoc (Bio-Rad). After detection of the signal, the membrane was stripped, blocked again, and incubated with the mouse anti-αtubulin antibody (Sigma-Aldrich) at a dilution of 1:500 for 1 h at RT. The membrane was washed and incubated with the antibody anti-mouse-HRP conjugate. The signal was detected by incubating with Pierce™ ECL Western Blotting Substrate (Thermo Scientific) for 5 minutes and using the ChemiDoc equipment (Bio-Rad).

### Immunofluorescence microscopy

BSF cells were maintained in HMI-9 (complete and Gln-free media), washed twice in PBS, and fixed with paraformaldehyde 2.4%, at 4 °C. After fixation, the cells were washed in PBS and resuspended in phosphate/glycine buffer. The cells were permeabilized by the addition of 0.1% Triton X-100, for 5 min and washed with PBS + 1% BSA. Then the cells were incubated with a 1:5,000 dilution of the antibody GT335 (AdipoGen) plus a 1:5,000 dilution of the antibody PolyE (anti-polyglutamate chain, AdipoGen) for 1 h at 4 °C. After incubation, the cells were washed twice with PBS+1% BSA. Then the parasites were incubated with 1:5,000 dilutions of each of the goat antibodies, anti-mouse IgG Alexa fluor™ 488 (Thermo Fisher Scientific) and anti-rabbit IgG Rhodamine Red™-X (Invitrogen), for 1 h at 4 °C, and protected from light. Cells were then washed once with PBS+1% BSA, twice with distilled water, transferred to slides, and covered with Fluromount G mounting medium (Invitrogen). Finally, cells were analyzed by confocal microscopy (Zeiss LSM780).

### Scanning Electron Microscopy (SEM)

BSF cells were maintained in the HMI-9 (complete or Gln-free media) for 24 h. After incubation, the cells (5 x 10^6^) were centrifuged (1,000 x *g*) and resuspended in Karnovsky’s fixing solution (2.5% glutaraldehyde, 4% paraformaldehyde in 0.1 M sodium cacodylate buffer, pH 7.2). The fixed cells were washed in a sodium cacodylate buffer and transferred to glass slides previously treated with poly-L-lysine. The samples were post-fixed by incubation in a 1% osmium tetroxide solution (sodium cacodylate buffer), washed, and dehydrated by increasing ethanol concentration solutions. The material was critical point dried and covered with a gold solution to be visualized by SEM (Jeol JSM6010PlusLA). The acquisition of images was performed at the Carlos Chagas Institute – FIOCRUZ, Paraná, Brazil.

### Transmission electron microscopy (TEM)

BSF cells (1 x 10^8^) were washed in TDB buffer and fixed for 1 h at 4 °C in a fixing solution (4% paraformaldehyde + 0.1% glutaraldehyde). After fixation, the cells were washed in a cacodylate buffer and incubated in a 1.5% osmium tetroxide solution (for 1 h at 4 °C). Cells were then washed (1,000 x *g*) once in the cacodylate buffer and three times with distilled water and incubated for 1 h in a 0.5% uranyl acetate solution at room temperature. Following a final wash step (1,000 x *g*) in distilled water, cells were dehydrated using increasing concentrations of ethanol (50%, 70%, 95%, and 100%) and propylene oxide. The EPON-embedding solution (2 g agar resin; 1 g DDSA; 1.08 g MNA; 0.04 g BBMA) was mixed (1/1) with propylene oxide, and the parasites were incubated in this solution for 1 h at RT. Cells were centrifuged again (1,000 x *g*), embedded in EPON (1 h at room temperature), collected by centrifugation (25 min at 1,000 x *g*), and transferred to the incubator for polymerization (12 h at 45 °C and 24 h at 60 °C). The specimens were sliced, contrasted by uranyl acetate (1 h at RT) and lead citrate (3 min at RT), and analyzed by TEM (Jeol JEM1400Plus) at the Carlos Chagas Institute – FIOCRUZ, Paraná, Brazil.

### Cryo-Soft X-Ray Tomography (Cryo-SXT)

#### Sample preparation

BSF cells (1 × 10^5^) were centrifuged (1,000 x *g*) and fixed in PBS + 2% paraformaldehyde. Aliquots of 3 μL were deposited onto Au-holey carbon Films for finder microscopy grids (Quantifoil R 2/2 on 200 *mesh gold finder grids*) with Au nanoparticles (BBI Solutions). The grids were blotted (Leica® EM GP Grid Plunger; Leica Microsystems), flash-frozen, and transferred into the full-field soft X-ray transmission microscope of the MISTRAL beamline at the ALBA Synchrotron (Barcelona, Spain) (43), where Cryo-SXT tomographic measurements of whole frozen hydrated cells were performed. The cryogenic conditions were maintained throughout the entire experiment.

#### Data acquisition and treatment

Tilt series at 520 eV X-ray energy were collected to enable 3D volume reconstruction of cells and their internal structures. At this energy, C exhibit high absorption relative to Oxygen, providing a strong natural contrast between C-rich membranes and the surrounding water-rich cytoplasm, without requiring staining or additional treatment after flash freezing. Given the relatively thin nature of the cells, a 25 nm Zone Plate lens was used as an optimal compromise between spatial resolution (approximately 30 nm half pitch) and depth of focus. The exposure time for each image was 1 s, and the typical angular range was ± 65 degrees, with one-degree increments. Each image of the tilt series was normalized by the incident intensity to obtain transmission values and then deconvoluted using a Wiener filter calculated from the lens PSF (44). To ensure accurate 3D reconstruction, the images were aligned along a common rotation axis, a step that is critical for reconstruction quality. The best alignment was achieved using the iterative algorithm implemented in the ARETOMO software package (45). The final aligned transmission tilt series were reconstructed with TOMO3D (46), using the SIRT iterative algorithm with 30 iterations. The cell volumes obtained were segmented using the Amira 3D software version 2021.2 (Thermo Fisher Scientific). The features of interest, including cell membranes, flagella, nuclei, nucleoli, mitochondria, and kinetoplasts, were manually highlighted and segmented. The segmentation was refined using the Magic Wand tool, and the surface of each segmented label was generated using the Generate Surface tool for surface visualization. Volume measurements were performed using the Label Analysis module of the AMIRA 3D software.

### Cell motility analysis

BSF motility was monitored using a sedimentation assay in a spectrophotometer (26,47). Briefly, the parasites were maintained in the standard HMI-9 (complete or Gln-free media) for 24 h. After the incubation period, 5 x 10^6^ cells of each culture were transferred to optical cuvettes. The density was monitored at 600 nm by using the spectrophotometer Evolution® 300 (Thermo Scientific), at 37 °C, for 2 h with 5 min intervals between readings.

### Glutamylproteome analysis

#### Enriched cytoskeleton fraction extraction

BSF cells were maintained in HMI-9 for 24 h. Thereafter, 8 x 10^7^ cells were washed twice (1,000 x *g*, 5 min, RT) and resuspended in PBS (500 µL). The enriched cytoskeleton fraction was extracted with 2 mL of 1% NP-40, PEME (48) (2 mM EGTA, 1 mM MgSO4, 0.1 mM EDTA, 0,1 M piperazine-N,N′-bis(2-acid ethanosulfonic)–NaOH (PIPES-NaOH), pH 6.9, 1 mM MgCl_2,_ and complete protease inhibitor cocktail (Sigma®) for 5 min at RT. The samples were centrifuged for 5 min, 5,000 x *g* at 4 °C, and the supernatant was removed. The complete cell lysis was confirmed by microscopy analysis. The flagellum was extracted with 1 mL of 1% NP40, PEME, 1 mM MgCl_2,_ pH 6.9, 1 M KCl, and a complete protease inhibitor cocktail, and incubated for 30 min at 4 °C. The samples were centrifuged for 5 min at 5,000 x *g* and 4 °C, resuspended in 4 volumes of cold acetone (-20 °C), and stored at -20 °C for 2 h. After thawing, samples were centrifuged (20 min, 16,000 x *g*, 4 °C), the supernatant discarded, and the pellet washed with 1 mL of cold acetone (5 min, 16,000 x *g*, 4 °C). Finally, the pellet was dried in a speed vac and analyzed by liquid chromatography-tandem mass spectrometry.

#### Liquid chromatography tandem mass spectrometry

Precipitated proteins were dissolved in 100 mM NH_4_HCO_3_ and digested with 0.5 µg of sequencing-grade trypsin overnight at 37 °C. Digested peptides were desalted in a C18 microspin column (Nested Group) following the manufacturer’s recommendations. Peptides were analyzed using a nanoAcquity UPLC (Waters) coupled to a Q-Exactive mass spectrometer (Thermo Fisher Scientific), as previously described (49). The data were analyzed using MaxQuant software v.1.6.17.0 by searching against the *T. brucei brucei* 927 sequences from Uniprot Knowledgebase. Searching parameters included: (1) tryptic cleavage sites in both termini with up to 2 missed cleavage sites and (2) methionine oxidation and glutamate glutamylation as variable modifications. The matching between runs function was enabled. The remaining parameters were used as the software default.

## Acknowledgments

The authors thank Dr. Ernesto Nakayasu from the Pacific Northwest National Laboratory for insightful discussions and critical reading of the manuscript. The authors would also like to acknowledge Dr. Mario Costa Cruz for acquiring confocal images at the Centro de Facilidades para Apóio à Pesquisa (CEFAP-USP), University of São Paulo, Brazil. The authors further thank the Confocal and Electron Microscopy Platform (RPT07C), Technological Platforms Network, Oswaldo Cruz Foundation (FIOCRUZ). Cryo-SXT measurements were performed at the MISTRAL beamline of ALBA Synchrotron, with the collaboration of ALBA staff, Cerdanyola del Vallès, Spain. The authors also acknowledge Jessica do Nascimento Faria de Souza for assistance with plunge-freezing of sample grids.

This work was supported by the Fundação de Amparo à Pesquisa do Estado de São Paulo (FAPESP) grant 2021/12938-0 (awarded to AMS) and the Conselho Nacional de Pesquisas Científicas e Tecnológicas (CNPq) grant 307487/2021-0 (awarded to AMS).

F.S.D. and S.M. were FAPESP fellows, and R.O.O.S. was a CNPq fellow during the development of this work. S.M. is currently a FAPESP fellow.

